# DNA hydroxymethylation reveals transcription regulation networks and prognostic signatures in multiple myeloma

**DOI:** 10.1101/806133

**Authors:** Jean-Baptiste Alberge, Florence Magrangeas, Mirko Wagner, Soline Denié, Catherine Guérin-Charbonnel, Loïc Campion, Michel Attal, Hervé Avet-Loiseau, Thomas Carell, Philippe Moreau, Stéphane Minvielle, Aurélien A. Sérandour

## Abstract

Multiple myeloma (MM) is a plasma cell malignancy that remains challenging to cure despite a substantially improving median survival. During the last decade, DNA copy number variation and gene expression studies have described the pathology and its heterogeneity among patients. Epigenetic modifications play important roles in MM, but they are rarely associated with clinical aspects of the disease. In this epigenomics study, we produced quantifications of genomic 5-methylcytosine (5mC) and of 5-hydroxymethylcytosine (5hmC) as well as genome-wide maps of hydroxymethylation to analyse myeloma cells taken from a cohort of 40 newly diagnosed and homogeneously treated patients. We found 5hmC to be globally depleted in MM compared to normal plasma cells, as well as being reduced in advanced clinical stages of the disease. From the hydroxymethylome data, we observed that remaining 5hmC is organised in large peak clusters and is associated with well-known disease-related genes. Based on their signal correlation, these 5hmC peak clusters can be gathered in 2 regulation networks involving core transcription factors such as *IRF4, MYC, PRDM1* and *TCF3*. By performing paired hydroxymethylomes at diagnosis and at relapse, we found the disease progression to be heterogeneous and patient-specific. We found that the location of 5hmC at tumor suppressor *TP53INP1* is associated with better outcome while global high level of 5hmC tends to be associated with better overall survival. Together, our study suggests that 5hmC provides new biological insights of the disease severity and progression, and can be used for retrospective studies.

**Key Points:**

- 5hmC is globally depleted in MM and even more in advanced stages of the disease.
- 5hmC is locally detected at transcriptionally active regions of MM where its presence can be associated with survival.

## Introduction

Multiple Myeloma (MM) is a plasma cell (PC) neoplasm with an incidence rate of 5/100,000 in Europe and accounts for approximatively 1 percent of all cancers. Median survival of patients has greatly improved in the last decade^1^ with the use of novel strategies such as autologous stem cell transplantation and new sets of drugs: immunomodulators, proteasome inhibitors, histone deacetylase (HDAC) inhibitors and monoclonal antibodies.^2^ Yet treatment remains challenging as nearly all patients ultimately relapse with the emergence of a resistant subpopulation of malignant plasma cells. Malignant clones show an heterogeneous range of mutations and chromosomal abnormalities along with heterogeneous chromatin and epigenetic dysregulations at diagnosis and at relapse that affects biological pathways (MAPK, NF-*κ*B, DNA-repair).^3^

Genomic and transcriptomic studies have allowed a better understanding of the disease and identified key transcription factors involved such as IRF4, MYC, PRDM1 and XBP1.^3–5^ Recent epigenomics technologies can help to deepen our knowledge of the transcriptional programs shaping MM. However epigenomics analysis through histone marks profiling with chromatin immunoprecipitation (ChIP-seq) or open chromatin mapping with Assay for Transposase-Accessible Chromatin (ATAC-seq) can be hard to set up on a cohort with limited material over a long range of time and thus give limited insights on the disease establishment and relapse. Studying DNA epigenetics marks is more adapted to this challenge.

Agirre and colleagues^6^ described DNA methylation in an important number of MM samples. They identified very heterogeneous levels of methylation from one patient to another. They found that despite a global hypomethylation, local and extensive hypermethylation is present in MM at intronic enhancer regions that are associated with stem cell development.

Chatonnet and colleagues^7^ recently identified several hydroxymethylated CpGs in MM samples with limited association to disease severity or prognosis, and to our knowledge, the genome-wide mapping of 5hmC (hydroxymethylome) has never been studied in a well-established cohort.

Oxydative states of 5-methylated Cytosine (5mC) on genomic DNA have been identified a decade ago^8–10^ : 5hmC, 5fC and 5caC. The TET proteins TET1/2/3 are responsible for the 5mC oxidation. 5fC and 5caC are almost undetectable in genomic DNA unless the glycosylase TDG gene is knocked-out^11, 12^, whereas 5hmC can be found in all cell types at various levels^13–15^. 5hmC is believed to be a DNA demethylation intermediate in a process involving TET proteins, TDG and the Base Excision Repair system.^16^ However 5hmC is also shown as a stable DNA modification.^17, 18^ Despite this discrepancy, 5hmC is commonly accepted as a DNA mark associated with active chromatin^19–21^ and is a powerful way to identify active genomic regions associated with a disease directly from genomic DNA or more recently from circulating DNA.^22, 23^

In this study, we characterized the 5hmC signal genome-wide on plasma cell DNA from 40 patients newly diagnosed with MM between 2010-2012 and representative of the main molecular subtypes, including 4 paired relapse samples, and of the plasma cells of 5 control individuals.

## Methods

### Normal plasma cells purification

After informed consent, the femoral canal of individuals with isolated hip osteoarthritis who were otherwise healthy was probed with a metal suction device following femoral neck removal. Bone marrow cells were suctioned into a tube that contained EDTA, placed on ice and immediately transported to our laboratory. BMMCs were purified by Ficoll. Normal plasma cells were FACS-sorted using a BD FACSAria III as CD38/CD138 positive and CD3/CD13/CD33 negative (antibodies from Becton Dickinson, ref. 345807, 555332, 555394, 555450 and 562935).

### Myeloma cells purification

Bone marrow biopsies were realized on MM newly diagnosed patients from cohort IFM-DFCI 2009 (Intergroupe Francophone du Myelome - Dana Farber Cancer Institute).^24, 25^ All patients signed an informed consent form approved by the Toulouse Ethics Committee. All included patients were newly diagnosed with symptomatic MM based on International Myeloma Working Group 2003 Diagnostic Criteria.^24^ All the samples were collected in France and processed at the University Hospital of Nantes. Bone Marrow Mononuclear Cells (BMMCs) were purified by Ficoll. Plasma cells were purified with anti-CD138 beads (Robosep platform, StemCell Technologies) and the CD138-positive percentage of cells was checked by immunofluorescence microscopy.

### Genomic DNA extraction

Genomic DNA was extracted with Qiagen Allprep DNA/RNA Mini Kit (ref. 80204). DNA samples were dosed by DNA HS QuBit and the absence of contaminant RNA checked by RNA HS Qubit.

### Patients selection for this study

In this study, we have selected 40 patient samples with: low level or absence of RNA in DNA samples, low level of rRNA in RNA-seq data, enough DNA material available (at least 1 ug) and high percentage of CD138+ cells (98% in average in this study).

### Digest of genomic DNA and subsequent LC-MS analysis

The genomic levels of methylcytosine (mC) and hydroxymethyl-cytosine (hmC) were quantified using a mass spectrometry-based stable isotope-dilution method.^26^ For each LC-MS-measurement (technical replicate), 70 ng of genomic DNA (gDNA) were digested to the nucleoside level using the Nucleoside Digestion Mix (ref. M0649S) from New England BioLabs. To this reason, a solution of 70 ng gDNA in 38 *µ*L of milliQ-water was prepared. As heavy-atom-labeled internal standards, 1.28 pmol D_3_-mC and 0.193 pmol D_2_ ^15^N_2_-hmC in 6 *µ*L of milliQ-water were added to the solution, followed by 5 *µ*L of the Nucleoside Digestion Mix Reaction Buffer (10x), and 1 *µ*L of the Nucleoside Digestion Mix. After incubation for 90 min at 37°C, the mixture was filtered using an AcroPrep Advance 96 filter plate 0.2 *µ*m Supor from Pall Life Sciences and subsequently analyzed by LC-MS. For each biological sample, two independent measurements (technical replicates) were performed. Quantitative LC-ESI-MS/MS analysis of the enzymatically digested DNA samples was performed using an Agilent 1290 UHPLC system coupled to an Agilent 6490 triple quadrupole mass spectrometer. The UHPLC-conditions used and the settings of the mass spectrometer were the same as previously published by Traube and colleagues.^26^

### Selective chemical labeling of 5hmC coupled with sequencing (5hmC-seq)

For each sample, 550 ng of genomic DNA was sonicated with a Bioruptor Pico in Tris 10 mM pH 8 to obtain DNA fragments of 300 bp in average. 25 pg of 5hmC control spike-in were added to the sonicated DNA (control provided by the kit HydroxyMethylCollector, Active Motif, ref. 55013). 50 ng of DNA was conserved at this stage to make the input library later. The remaining DNA was processed using the HydroxyMethylCollector kit (method from Song and colleagues^27^) to glycosylate and biotinylate specifically the genomic 5hmC. After glycosylation and biotinylation the DNA was purified with Ampure beads (Beckman Coulter, ref. A63881). The DNA fragments containing the biot-glu-5hmC were purified with Streptavidine beads (Active Motif, ref. 55013), eluted, purified with Ampure beads and finally eluted in 50 uL Tris pH 8. The 5hmC-seq libraries were prepared with the kit NEBNext Ultra II DNA library prep kit for Illumina (ref. E7645S) and indexed with NEBNext dual indexed primers (E7600S). The libraries were quality-checked by HS DNA Agilent BioAnalyzer (Supplementary Figure 1), dosed by DNA HS Qubit, pooled and submitted to the genome sequencing platform for Single-Read 50 bp Illumina HiSeq-2500 Rapid Run sequencing.

### RNA-seq librairies

As the IFM-DFCI did not include RNA-seq data from patients at relapse, we produced RNA-seq data at diagnosis and at relapse on 4 selected MM patients (patients number MM 2, MM 5, MM 7 and MM 21). The RNA-seq libraries were prepared using the NEBNext Poly(A) mRNA Magnetic Isolation Module (NEB, ref. E74905) and the NEBNext Ultra II Directional RNA Library Prep Kit for Illumina (NEB, ref. E7760S) and sequenced by an Illumina Rapid Run HiSeq 2500 Single-Read 50 bp.

### Bioinformatics

#### 5hmC-seq analysis

5hmC-seq libraries were trimmed with Cutadapt v1.18^28^ and mapped to human reference genome GRCh38 using Bowtie2 v2.2.8^29^ with default parameters. Duplicates and reads of low mapping quality (*Q <* 30) were discarded. 5hmC peak clusters were defined similar to Boeva and colleagues^30, 31^ with a stitching distance of 12.5kb (Supplementary Figure 2). Blacklisted regions consist of chromosomes X, Y, ENCODE reference blacklist and peaks called on input-seq libraries.

Genes and 5hmC peaks clusters were associated to the Topologically Associated Domain (TAD) of the B-cell derived cell line GM12878 they are located in similar to Boeva and colleagues.^31^

Group specific 5hmC peaks clusters were identified by computing the average Log2 Fold-Change signal per region between groups and tested with a Wilcoxon rank-sum test adjusted with a Benjamini-Hochberg correction. For paired samples (two replicates of each diagnosis and relapse), we used DiffBind^32^ with default parameters to find differentially enriched 5hmC regions.

Potential transcription factor binding sites and core regulatory circuiteries were found with CRCMapper and MEME.^33, 34^

Network clustering of 5hmC was computed on the correlation matrix of peak clusters in the cohort. A graph adjacency matrix was calculated from the correlation between peak clusters with a threshold of 0.715. Every correlation above 0.715 is counted as an adjacency between two peak clusters. This value was chosen to limit the number of non-connected components in the graph and to keep about 400 peak clusters in the analysis. The two sub-networks were then identified from the hierarchical clustering (Ward’s method) of the first two Eigen components of the symmetric normalized Laplacian matrix.^35^

Bioinformatics code is hosted at: https://gitlab.univ-nantes.fr/alberge/5hmc-myeloma-analysis.

#### RNA-seq analysis

IFM-DFCI RNA-seq data were obtained from Cleynen and colleagues^36^ or sequenced by us. RNA-seq libraries were trimmed using Cutadapt v1.13^28^ with parameter − *m* 20 and aligned to hg38/GRCh38 using STAR v2.5.3a^37^ with default parameters. Genes were quantified using featureCounts^38^ from the R package Rsubread and the Gencode v28 genome annotation. RefSeq genome annotation was used in parallel as a reference for transcription factor selection.

#### ChIP-seq and ATAC-seq analysis

ChIP-seq and ATAC-seq libraries were downloaded from the European Nucleotide Archive (project number PR-JEB25605^39^) and treated with the pyflow-ChIPseq^40^ and pyflow-ATACseq pipelines respectively with default parameters. Chromatine-states genome annotation was realized using ChromHMM^41^ on ChIP-seq data from MM.1S cell line.

#### Survival analysis

Time-to-event was calculated from the randomization to the event date, i.e. death for OS, death or relapse for PFS, or to the last follow-up date.

For MS quantitative variables, cohorts were split at the median value of high and low 5hmc (and 5mC). Survival curves were calculated using the Kaplan-Meier method and groups were compared using a Log-rank test.

For 5hmC-seq data, at univariate step, SAM^42^ was run to extract the 5hmC regions that are significantly correlated with overall survival (*q*_*value*_ *<* 0.05) with 100 permutations and Δ = 0.2. For multiple regions testing (scores SC1 and SC2), the cohort was split at median value of 5hmC enrichment. 5hmC of each patient was summarized by the weighted mean of independently significant parameters. Variables were standardized and weights were based on multivariate Cox’s coefficients changed to +1 or −1 according to their sign to avoid overfitting. Survival curves were calculated using the Kaplan-Meier method and groups were compared using a Log-rank test.

### Data Sharing Statement

Sequencing data are accessible at ENA under accession PRJEB32800. Mass spectrometry data are available in Supplemental Table.

## Results

### The 5hmC regions landscape in MM

We studied a cohort of 40 patients newly diagnosed with MM between 2010-2012 and 5 healthy bone marrow donors (Figure 1A).

**Figure 1:**
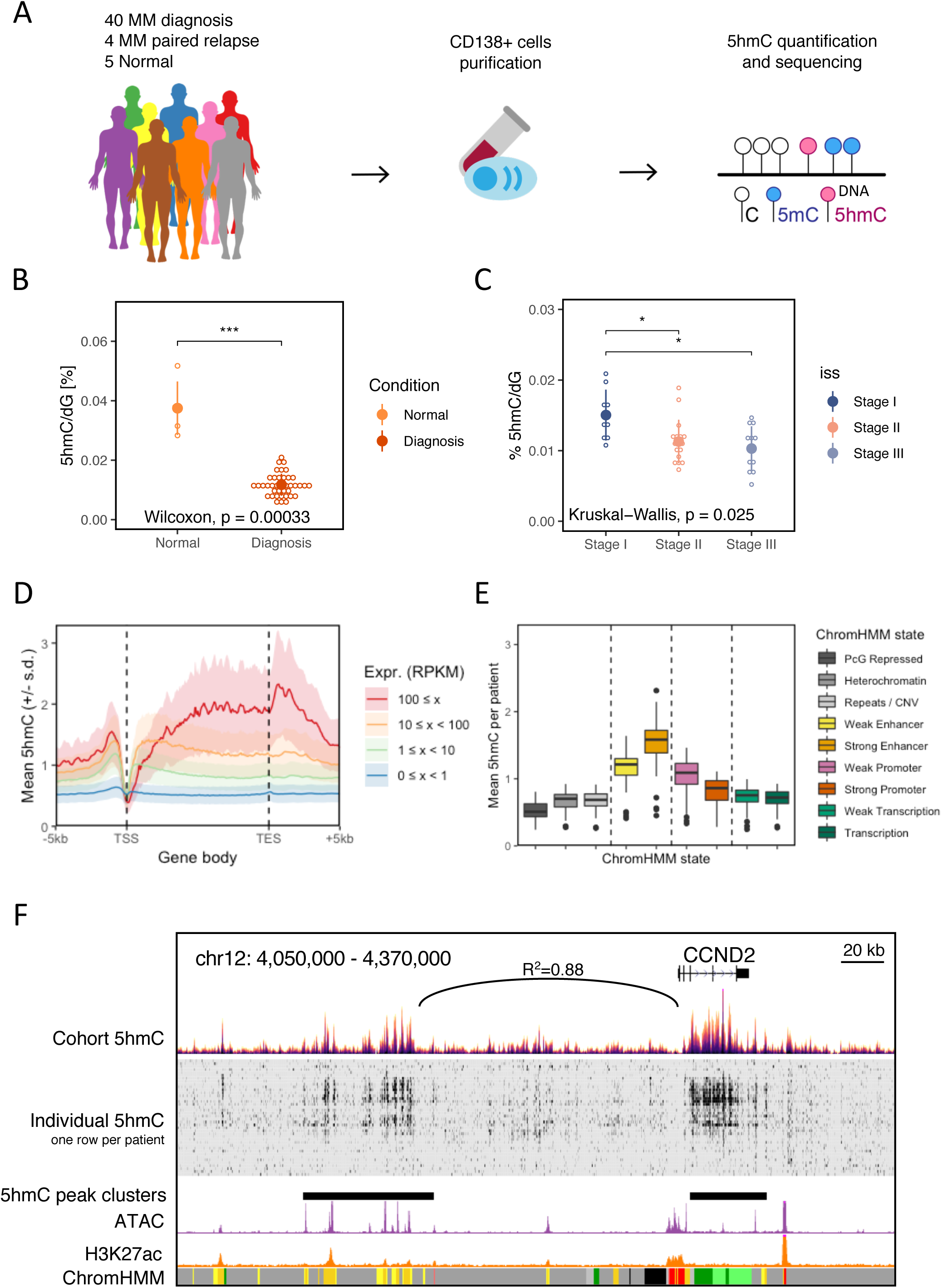
5hmC is associated with patient classification and is located at expressed genes and at strong enhancers. (A) Experimental pipeline leading to MS and 5hmC-seq analysis in plasma cells from patient bone marrows. (B) Dot plot of 5hmC global quantification by MS in normal plasma cells from healthy donors (N=5), and of myeloma cells of patients at diagnosis (N=39). (C) Dot plot of 5hmC global quantification by MS by disease stage (ISS I N=9; ISS II N=17; ISS III N=12; NA=1). ISS, International Staging System. (D) Average 5hmC signal at genes across the whole 5hmC dataset. The average 5hmC signal is plotted by class of gene expression level in RPKM. (E) Distribution of the 5hmC-enriched regions at the different ChromHMM chromatin states across the 5hmC dataset. (F) 5hmC, ATAC, H3K27ac signal and ChromHMM states at the *CCND2* genomic locus. The 5hmC signal correlation between the *CCND2* gene and its putative enhancer is indicated. ATAC and H3K27ac data were published by Jin and colleagues.^39^

We quantified by MS the global level of 5mC and 5hmC in our 49 samples (40 diagnosis in 5mC, 39 in 5hmC, 4 relapses and 5 normal plasma cell samples). We found the 5mC and the 5hmC to be both significantly reduced in MM compared to normal plasma cells (NPC; Figure 1B and Supplementary Figure 3A). The amount of 5hmC is reduced in average by about 4 fold between NPC and MM (p_value_*<*0.001). We found that 5hmC, and not 5mC, is reduced in MM Stage II/III compared to Stage I regarding to the International Staging System,^43^ a classification of patients based on *β*_2_-microglobulin and albumin levels with a strong prognosis value (p_value_=0.025, Figure 1C and Supplementary Figure 3B). 5mC and 5hmC global levels were not correlated to the sex or the age of the patients (Supplementary Figure 3C-3F).

These results encouraged us to characterize the hydroxymethylome by 5hmC-seq^27^ to identify the residual genomic regions marked by 5hmC in MM.

We confirm that the level of 5hmC signal in enriched at gene bodies and is positively associated with RNA expression as expected from previous studies^20, 27, 44–46^ (Figure 1D).

The 5hmC signal is particularly enriched at strong enhancers and to a lesser extend at weak enhancers (Figure 1E) when we used a functional annotation from the MM.1S cell line.

Inspection of regions of interest showed that strong levels of 5hmC can be associated between distal loci (Figure 1F). For instance at the *CCND2* locus, we found a strong correlation (*R*^2^ = 0.88) between 5hmC signal in the gene body (chr12:4,278,700-4,312,900) and 5hmC signal at an extragenic centromeric region located 120 kb upstream (chr12:4,106,500-4,164,700; Supplementary Figure 4A). Both 5hmC signals correlate well with *CCND2* RNA expression and both regions are located in the same topogical domain according to HiC data^47, 48^ (chr12:3,850,000-4,800,000; Supplementary Figure 4B). The upstream region is also marked by ATAC and H3K27ac signals in MM patients^39^ suggesting that this genomic region is functionally active. This strongly suggests that this upstream region is an enhancer of *CCND2* gene.

Hence we pursued the analyses using 1816 5hmC peak clusters. We determined the 5hmC peak clusters common to at least two patients and performed an unsupervised Principal Component Analysis (PCA). PCA plot shows a great heterogeneity of MM compared with Normal patients (Figure 2A). PC1 is driven by *CCND1* (x-axis, left-side) versus *CCND2* (x-axis, right-side). PC2 is driven by *CCND1* (y-axis, bottom) versus *FGFR3* -*MMSET* (y-axis, top) (Supplementary Table), the three loci being of major interest in MM. Patients samples were then classified in 5 groups: MMSET (translocation t(4;14); 9 patients), CCND1 (RNA expression over 800 Transcripts per Million; 11 patients), hyperdiploid (16 patients, at least 2 odd chromosomal gains), others (MM in none of the aforementioned groups; 4 patients) and normal plasma cells (Normal; 5 donors).

**Figure 2:**
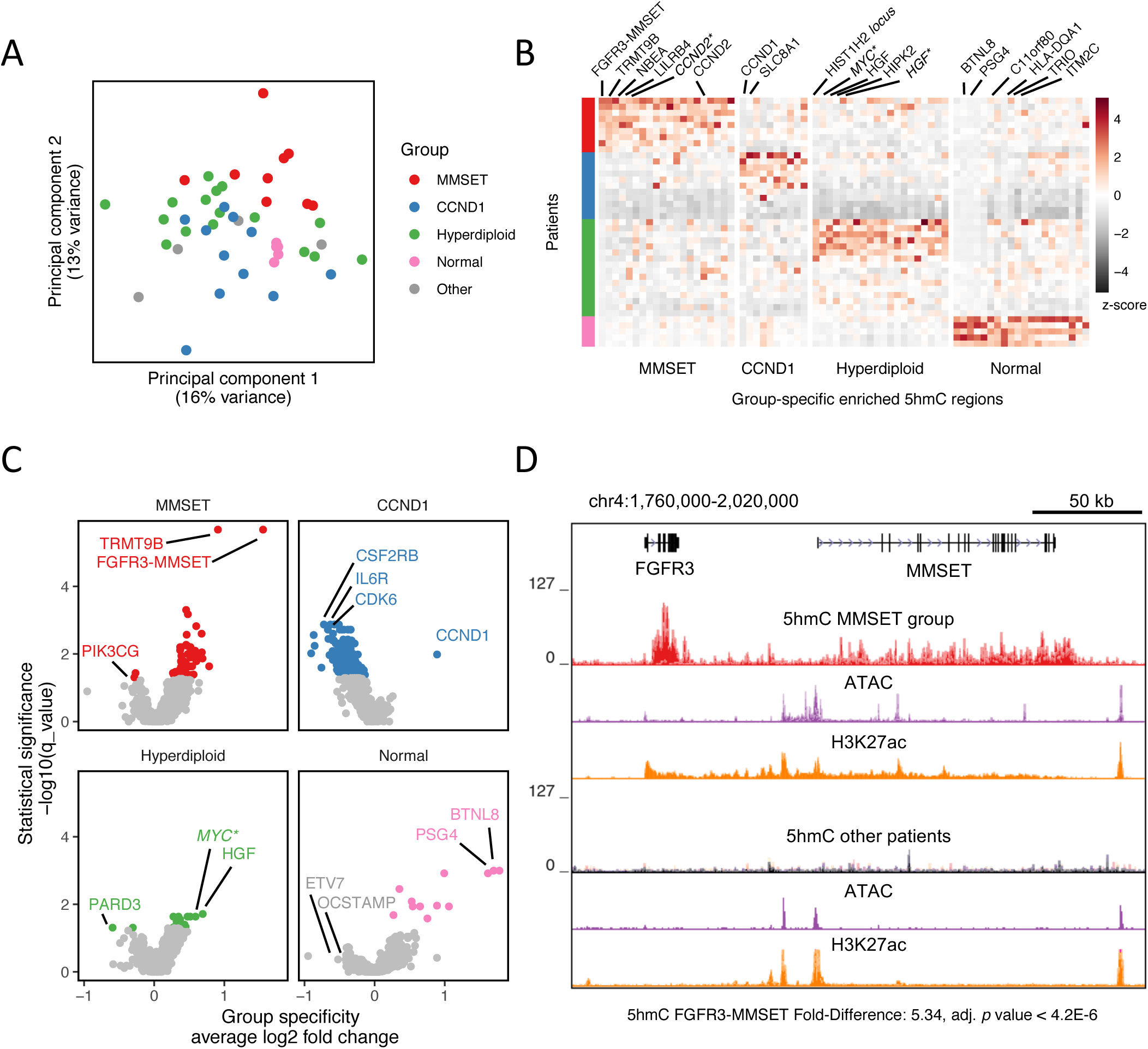
5hmC peak cluster are located at major functional genes in MM and can be associated with specific clinical subgroups. (A) 5hmC signal PCA plot showing the 40 MM samples (groups MMSET, CCND1, Hyperdiploid and Other) and the 5 normal plasma cells (group Normal). (B) Heatmap showing the 5hmC signal at the most differential 5hmC peak clusters between the groups MMSET, CCND1, Hyperdiploid and Normal. Asterisks stand for proximal non-genic loci. (C) Plot showing the 5hmC peak clusters that are specifically enriched in patient groups MMSET, CCND1, Hyperdiploid and Normal. Significant regions are colored. (D) 5hmC, ATAC and H3K27ac signals at the *FGFR3-MMSET* locus in the MMSET patient group and the other patients.

For each group, we determined specific 5hmC peak clusters (Figures 2B, 2C, Supplementary Figure 5A and Supplementary Table). Remarkably, the strongest specific 5hmC peak cluster for the group MMSET is the locus *FGFR3* -*MMSET* (p_value_=1.6 10^−6^) followed by *CCND2, LILRB4, NBEA* and *TRMT9B* (Figure 2D and Supplementary Figure 5B). This strong 5hmC enrichment in MMSET patients is also associated with strong H3K27ac and ATAC-seq signals^39^ compared to the other MM patients. This result suggests that the translocation between the *FGFR3* - *MMSET* locus and the *IgH* locus is associated massive 5mC oxidation together with chromatin opening and very high transcription level (Figure 2D).

Remarkably, in patients from the CCND1 group, the strongest and most specific 5hmC peak cluster was located in the *CCND1* gene itself (Figure 2C and Supplementary Figures 5C, p_value_=0.05). Again this suggests that the translocation event between *CCND1* and *IgH* or the novel overexpression of *CCND1* induces very high 5mC oxidation together with a strong *CCND1* transcription activity even though the classification of CCND1 patients was done based on RNA level in absence of FISH data.

The hyperdiploid group shows strongly specific 5hmC signal at *HGF* (Hepatocyte growth factor, p_value_=0.02) and at the locus of *MYC* oncogene (Figure 2B, 2C, Supplementary Figure 5D, p_value_=0.02).

Normal plasma cells are enriched in 5hmC at *BTNL8, C11orf80, ITM2C, PSG4* and *TRIO* genes (Figure 2B, 2C). We found high 5hmC signal together with H3K27ac and ATAC signals^39^ at several important genes implicated in MM including *CCND1, FGFR3* -*MMSET, IRF4, PRDM1* and *XBP1* (Supplementary Figures 6A-F). Specifically, H3K27ac super-enhancers from a patient with t(4;14), del13 and amp1q from Jin and colleagues^39^ and the 5hmC peak clusters found in one of our patients with the same genomic alterations are located at similar genomic loci *CREB3L2, FNDC3B, MMSET, ST6GAL1* and *ZBTB38* (Supplementary Figures 5E,F).

### Transcriptional regulatory networks in MM

To uncover transcriptional regulation networks shaping multiple myeloma, we next measured the signal correlation between the 1816 5hmC peak clusters among the 40 patients and searched for correlation networks. A correlation matrix of 407 5hmC peak clusters was obtained, in which 2 correlation networks (network 1 and network 2, Figure 3A) were defined through hierarchical clustering (of resp. 242 and 165 5hmC peak clusters). To find the core transcription factors (TF) involved in these 2 networks, we did a core transcriptional regulatory circuits analysis^34^ for each patient based on 5hmC and RNA expression data. We found that core TFs motifs were differently enriched between the 2 networks. Network 1 was enriched with TFs motifs such as KLF6/13, MYC, SP3, SREBF2, TCF3, USF2 and ZBTB7A, while network 2 was enriched with ARID3A, ATF4, FOXO3, IRF4/7, NFIL3 and PRDM1 motifs (Figures 3B, 3C, 3E). The ten most expressed TF were ATF4, KLF6/13, IRF1/4, PRDM1, TCF3, USF1/2 and XBP1 (Supplementary Figure 7A). These core transcription factors are shared between the two networks (Supplementary Figure 7B). For each patient, we averaged 5hmC signal in networks 1 and 2 and found the patients to be continuously distributed in terms of network scores rather than being clustered in one of the networks (Figure 3D). Through GREAT analysis,^49^ network 1 was shown to be associated with negative regulation of cytokine production and negative regulation of tumor necrosis factor production, while the network 2 was associated with immune response-regulating cell surface receptor signaling pathway and immune response-regulating signaling pathway (Figure 3F). Network analysis suggests that MM could be shaped by 2 regulation networks associated with specific transcription factors with partially redundant functions.

**Figure 3:**
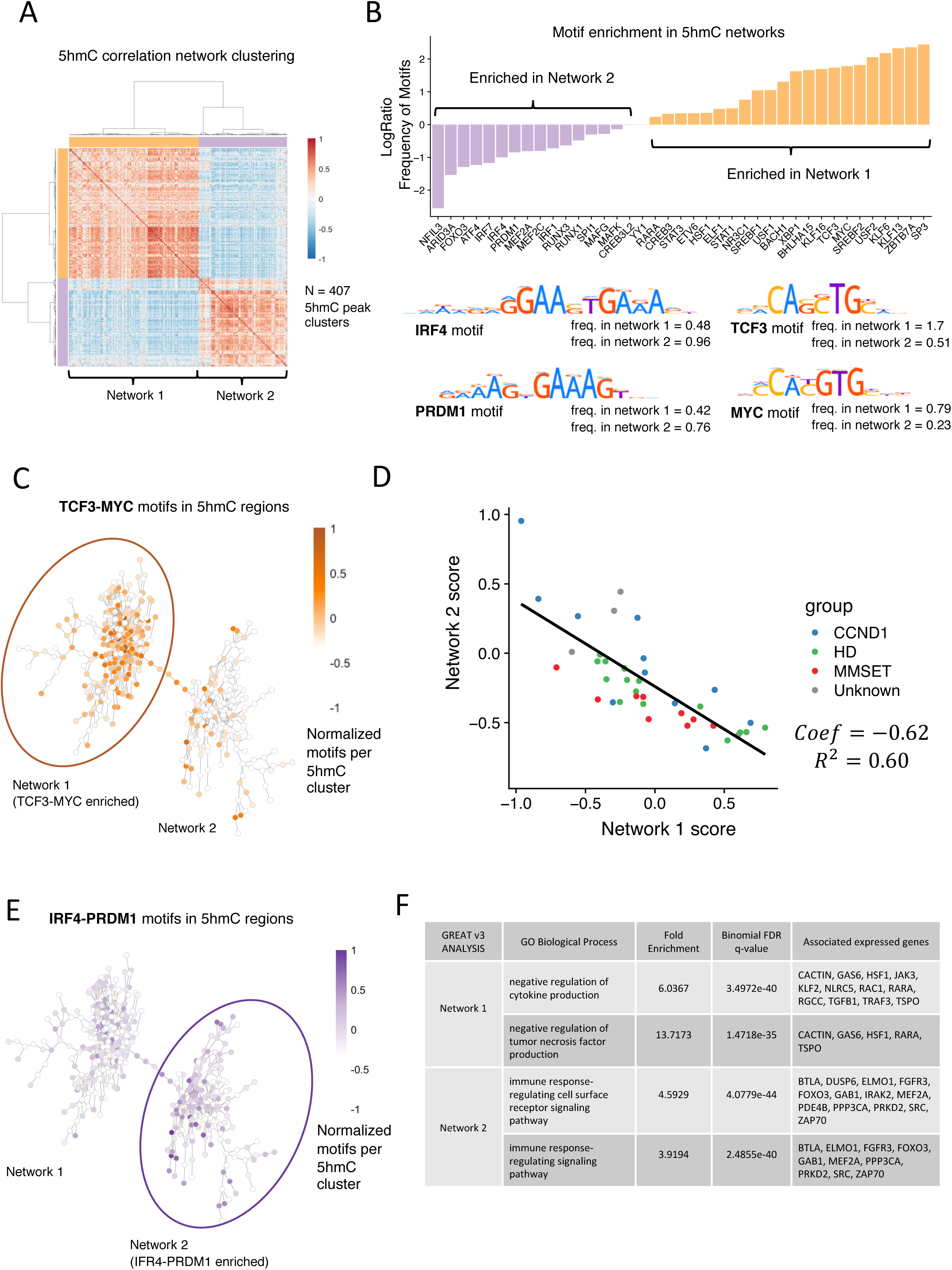
5hmC peak clusters are divided in two anti-correlated regulation networks. (A) Heatmap representing the signal correlation between the 5hmC peak clusters (407 correlated out of 1816). Clustering isolates two sets of regions that we called network 1 and network 2. (B) Motif enrichment of the core transcription factors in network 1 and 2 identified from the core transcriptional regulatory circuits analysis. (C) Network representation of the 5hmC peak clusters linked with the strongest correlations. Each dot is a 5hmC peak cluster. Edges are represented if the correlation between 2 nodes is greater than 0.715. Color intensity represents the normalized number of motifs TCF3-MYC in the 5hmC peak clusters. (D) Scoring of patient samples in both networks 1 and 2 with linear regression. (E) IRF4-PRDM1 motif occurences in the networks representation. (F) Gene ontologies predicted by GREAT^49^ for the networks 1 and 2.

### 5hmC dynamics between diagnosis and relapse in MM

To identify the genomic regions associated with MM progression, we next mapped 5hmC in 4 MM pairs (diagnosis and first relapse) and identified differentially hydroxymethylated regions with two replicates for each condition (Figure 4A). CNV microarray data^50, 51^ and RNA-seq were used at both time points to assess the progression of each patient (Figure 4B).

**Figure 4:**
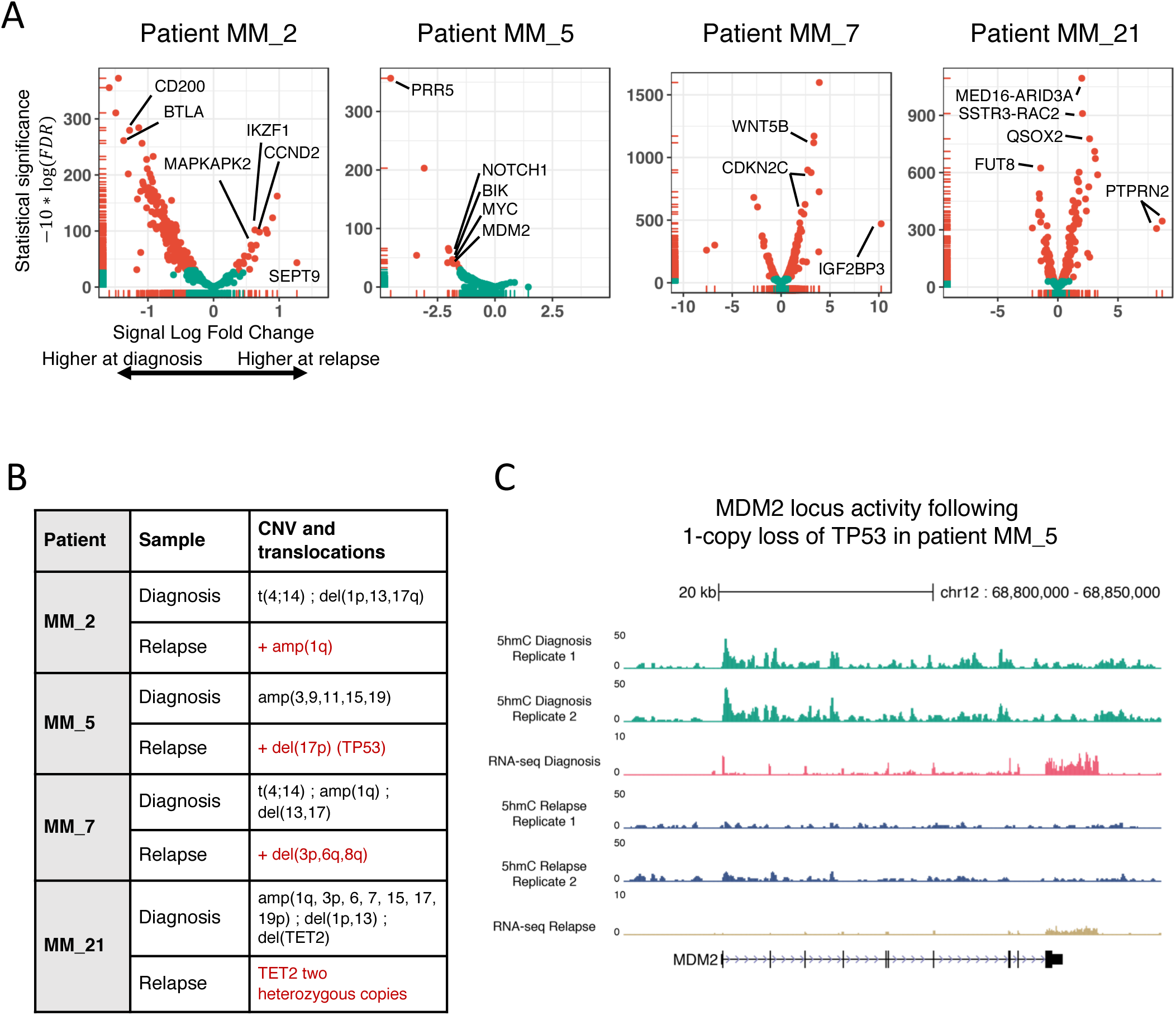
5hmC is dynamic and heterogeneous in MM between diagnosis and relapse. (A) Differential signal analysis of 5hmC peak clusters between diagnosis and relapse in patients MM 2, MM 5, MM 7 and MM 21. (B) Copy number variation history of MM patients between diagnosis and relapse. (C) 5hmC and RNA expression levels of the MDM2 gene at diagnosis and relapse in patient MM 5. At relapse, this patient acquires the deletion of the chromosome arm 17p (including a single-copy deletion of TP53).

The plasma cells of patient MM 2 showed a translocation of the *MMSET* locus (t(4;14)) with a deletion of chromosomes 1p, 13 and 17p at diagnosis. This subject progressed in 18 months, with his plasma cells displaying a third copy of 1q. Out of 560 consensus 5hmC regions, 269 (48%) are significantly reduced at relapse compared to diagnosis (FDR*<*0.05) while 20 (3.5%) are enriched at relapse, suggesting a global loss of 5hmC. Interestingly, we noted a significant gain of 5hmC at *CCND2* and *IKZF1* genes bodies at relapse (respectively 1,6 and 1,5 fold) associated with a higher transcriptional activity (respectively 2,4 and 1,5 fold). The drug lenalidomide induces IKZF1 proteasomal degradation,^52, 53^ suggesting that the up-regulation of *IKZF1*, together with an increase in *CCND2* expression could favor disease progression. We also noticed a gain of 5hmC at *MAPKAPK2/MK2* at relapse. This gene is located on 1q, which is gained at relapse by this patient, and has been recently described as a poor prognosis factor.^54^

At diagnosis, patient MM 5 displayed a classical hyperdiploid profile with amplification of chromosomes 3, 9, 11, 15, 19, where 675 regions were found to harbor 5hmC peak clusters. At relapse 24 months later, cells displayed a 1-copy loss of 17p, with 22 regions (3.2%) significantly decreased. Accordingly to *TP53* loss, expression and 5hmC signal of p53 target genes such as *MDM2* was found decreased (Figure 4C).^55, 56^

Patient MM 7 is another *MMSET* translocated patient (t(4;14)) with amplification of chromosome arm 1q and deletion of chromosomes 13 and 17p (Figure 4B) gaining deletions at relapse (3p, 6q, 8q). At relapse, 5hmC increases at several genes such as *CDKN2C, IGF2BP3* and *WNT5B* as well as their RNA expression (log fold-change *>* 3).

Patient MM 21 plasma cells were found to have a narrow *TET2* deletion at diagnosis, but surprisingly 2 heterozygous copies at relapse. TET2, which converts 5mC to 5hmC, is frequently mutated in cancer and especially in myeloid malignancies,^57^ but not in MM.^4^ 385 regions were found to harbour 5hmC, out of which 86 (22%) decreased and 117 (30%) increased at relapse. Global level of 5hmC decreased moderately despite the reappearance of TET2 (Supplementary Table).

### 5hmC and survival in MM

Survival analysis was first performed on the global level of 5mC and 5hmC. 5hmC (indicated as 5hmC/dG [%]) tends to be associated with Overall Survival (OS) although statistically non-significant (Hazard ratio (HR) = 2.6, CI=[0.9,7.8], p_value_=0.066) (Figure 5A). Yet 5hmC is less likely to be associated with Progression-Free Survival (PFS; Figure 5B; p_value_=0.3). 5mC global level does not show significant association with OS and PFS (Supplementary Figures 8A and 8B).

**Figure 5:**
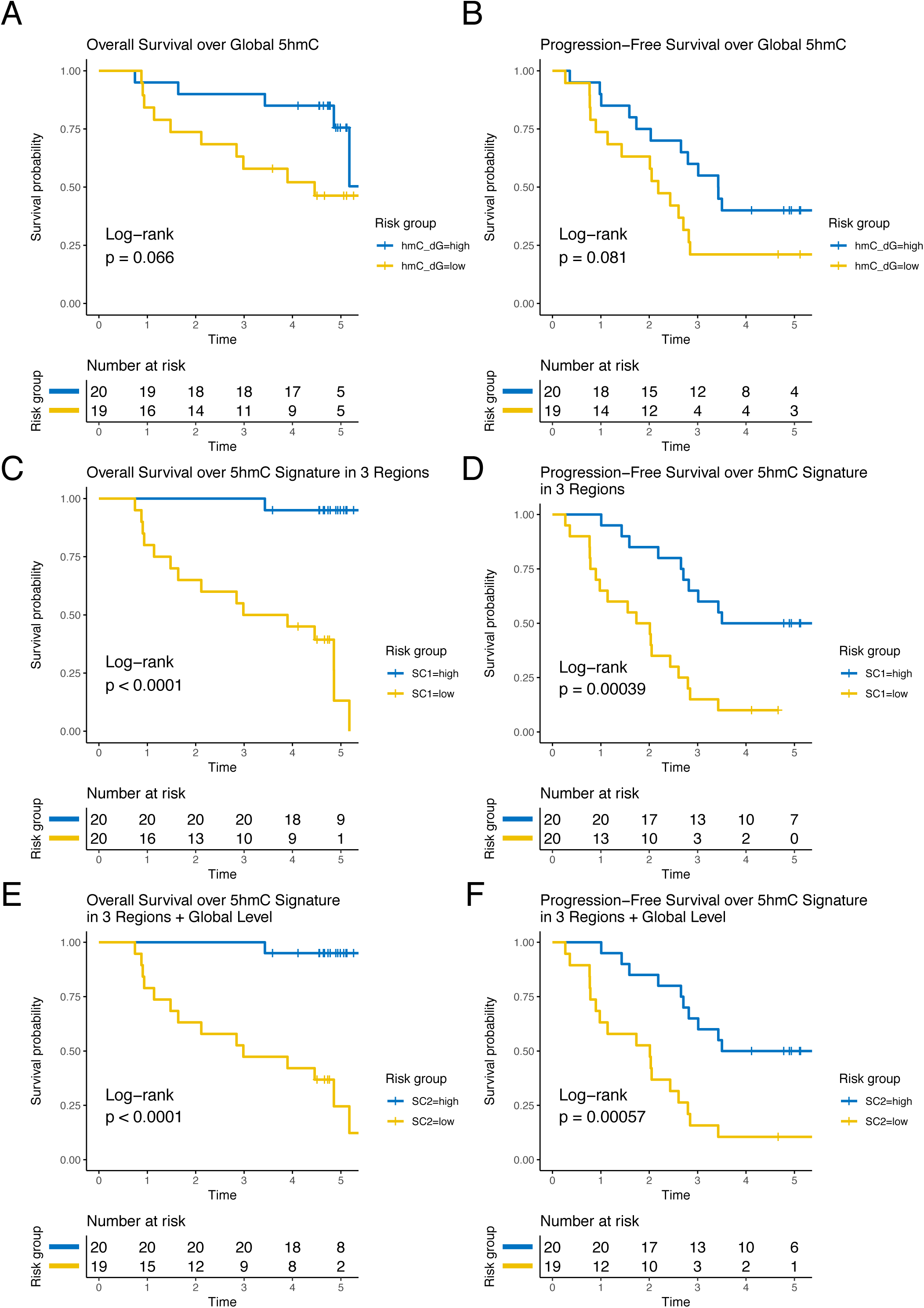
Genomic 5hmC levels are associated with Overall and Progression-Free Survival. Univariate and multivariate Overall Survival (OS) and Progression-Free Survival (PFS) analysis. 5hmC enrichment is split at the median value (for univariate) or median value of weighted mean (for multivariate, see Methods). (A) OS over global level of 5hmC in MS. (B) PFS over global level of 5hmC in MS. (C) OS over 5hmC enrichment at *TP53INP1, UPF1* and *RNF144A* genomic loci. (D) PFS over 5hmC enrichment at *TP53INP1, UPF1* and *RNF144A* genomic loci. (E) OS combining global level of 5hmC in MS and 5hmC enrichment at *TP53INP1, UPF1* and *RNF144A* genomic loci. (F) PFS combining global level of 5hmC in MS and 5hmC enrichment at *TP53INP1, UPF1* and *RNF144A* genomic loci. (Time: Number of years).

Three 5hmC genomic loci were found to separate independently the patients in 2 groups of good and poor survival. High 5hmC signal at the *TP53INP1* gene is associated with longer survival (HR=3.7, CI = [1.2-11], p_value_=0.017; Supplementary Figure 8C) and low 5hmC signal at *RNF144A* gene (or *UPF1* gene) is associated with poorer survival (data not shown).

The 3 independent 5hmC regions (*RNF144A, TP53INP1* and *UPF1*) were then summarized into a score SC1 (see Methods) that separates the cohort into 2 median groups of 20 patients of bad and good outcomes (HR=12, CI=[4, 244], p_value_=1 10^−6^ for OS; HR=3.8, CI = [1.7, 8.6], p_value_=4 10^−4^ for PFS; Figures 5C and 5D). The lower risk group shows higher level of 5hmC at *TP53INP1* gene and lower levels at *RNF144A* and *UPF1* genes. When applied to gene expression of *RNF144A TP53INP1* and *UPF1*, no significant impact on survival was found (HR=1.4, CI = [0.27, 2.0], p_value_=0.5; Supplementary Figure 8D).

We found that 5hmC global level is independent of the 5hmC enrichment at the 3 genomic loci. We combined the 4 variables into a second score SC2 (see Methods). Patients with better outcome show high global level of 5hmC, high 5hmC signal at *TP53INP1* and low 5hmC signals at *RNF144A* and *UPF1* (HR=24, CI = [3.2, 187], p_value_=7 10^−6^ for OS; HR=3.8, CI = [1.7, 8.4], p_value_=6 10^−4^ for PFS; Figures 5E and 5F).

## Discussion

We have quantified the epigenetic marks 5mC and 5hmC and mapped 5hmC in plasma cell DNA samples from 40 newly diagnosed MM patients and 5 donors. The global levels of 5mC and 5hmC are lower in MM compared to NPC. 5hmC is higher in Stage I MM compared to Stage II and Stage III patients. 5hmC tends to be positively associated with survival. These observations show that 5hmC is a marker of disease progression and severity.

Despite a global hypohydroxymethylation in MM, there still remain 5hmC marks at transcriptionally active regions. We revealed that 5hmC is strongly associated with active enhancers and identified active genomic regions specific to known MM molecular groups. We also found that H3K27ac-based super-enhancers from Jin and colleagues^39^ share similarities with our 5hmC peak clusters. These observations confirm that 5hmC is a valuable mark to study active genomic regions.

An unsupervised analysis shows that 5hmC at major oncogenes (*CCND1, CCND2, FGFR3* and *MMSET*) can separate hydroxymethylomes of MM patients, consistent with the existing classifications based on genetic abnormalities. 5hmC signals at the loci *CCND1* and *FGFR3* -*MMSET* are strong markers of the CCND1 and MMSET molecular groups respectively. This suggests that the translocation events are the major factors shaping MM hydroxymethylome. We discovered 2 networks of highly correlated 5hmC regions that offer a new prism to view MM transcriptional regulation. Our analysis connects target genes and regulatory regions to transcription factors. The 2 networks are predicted to be controlled by key transcription factors of well-known importance in MM (IRF4, MYC, PRDM1, and XBP1) as well as novel genes associated with the disease regulation (ATF4, IRF7, and TCF3). Of note, several MM-related transcription factors are under investigation to be drug-targeted and represent promising therapeutic targets.^58^ The gradual distribution of 5hmC signal between the two networks suggests that they may have redundant functions.

Between diagnosis and relapse, we found a highly dynamic and patient-specific distribution of the 5hmC signal. This reflects that MM progression is highly heterogeneous, although we could find consistency between 5hmC changes, expression and copy number variations.

We found 3 5hmC regions that were independently associated with overall survival. A high 5hmC signal at the *TP53INP1* gene is associated with a longer survival while low 5hmC signals at *RNF144A* and *UPF1* genes are associated with good outcome. TP53INP1 protein is shown as a tumor suppressor with antiproliferative and proapoptotic functions.^59, 60^ Interestingly, the association of TP53INP1 and overall survival is independent of del17p status.

Our study shows that the epigenetic mark 5hmC is valuable to discover active regulatory regions in genomic DNA from a cohort of patients without the need of chromatin extraction. It has been recently shown that it is also possible to map 5hmC on circulating DNA.^22, 23^ This makes 5hmC not only biologically valuable but also technically easier than ChIP-seq against histone marks or transcription factors. Taken together, these results show the value of epigenomics in retrospective studies and bring to light potential drug targets that drive the malignant transcriptome.

## Acknowledgments

We thank G. Salbert for the critical reading of this manuscript. We thank M. Devic, E. Douillard, E. Ollivier and N. Roi for their great technical support. We thank the Biogenouest sequencing platform GenoBird from Nantes for the Illumina sequencing. We thank the medical staff of the Nouvelles Cliniques Nantaises - Le Confluent that provided the bone marrow cells collected during hip replacement surgery.

This study was supported by the Chaire Mixte INSERM - Ecole Centrale de Nantes, the Intergroupe Francophone du Myelome, the Ligue Contre le Cancer, the I-SITE NexT (ANR-16-IDEX-0007) and the SIRIC ILIAD (INCa-DGOS-Inserm-12558). AJ was supported by a PhD Fellowship from INSERM and Région Pays de Loire. Research of M.W. and T.C. was supported by the Deutsche Forschungsgemeinschaft (SFB1309: TP A04, SFB1361: TP 02) and the European Research Council (ERC-AG) under the European Union’s Horizon 2020 research and innovation program (Project ID: EpiR: 741912).

## Authorship Contributions

J.-B.A., F.M., S.M, and A.S designed the project and analyzed the data. M.A., H.A.-L., P.M and S.M. created the cohort and provided patient samples. A.S. and S.D. produced the 5hmC-seq data. M.W. and T.C. performed the quantitative mass spectrometry measurements and interpreted the data. J.-B.A., C.G. and L.C. analyzed the survival data. A.S. and J.-B.A. wrote the manuscript. All authors read and corrected the manuscript.

## Supplementary Figure Legends

**Supplementary Figure 1:**
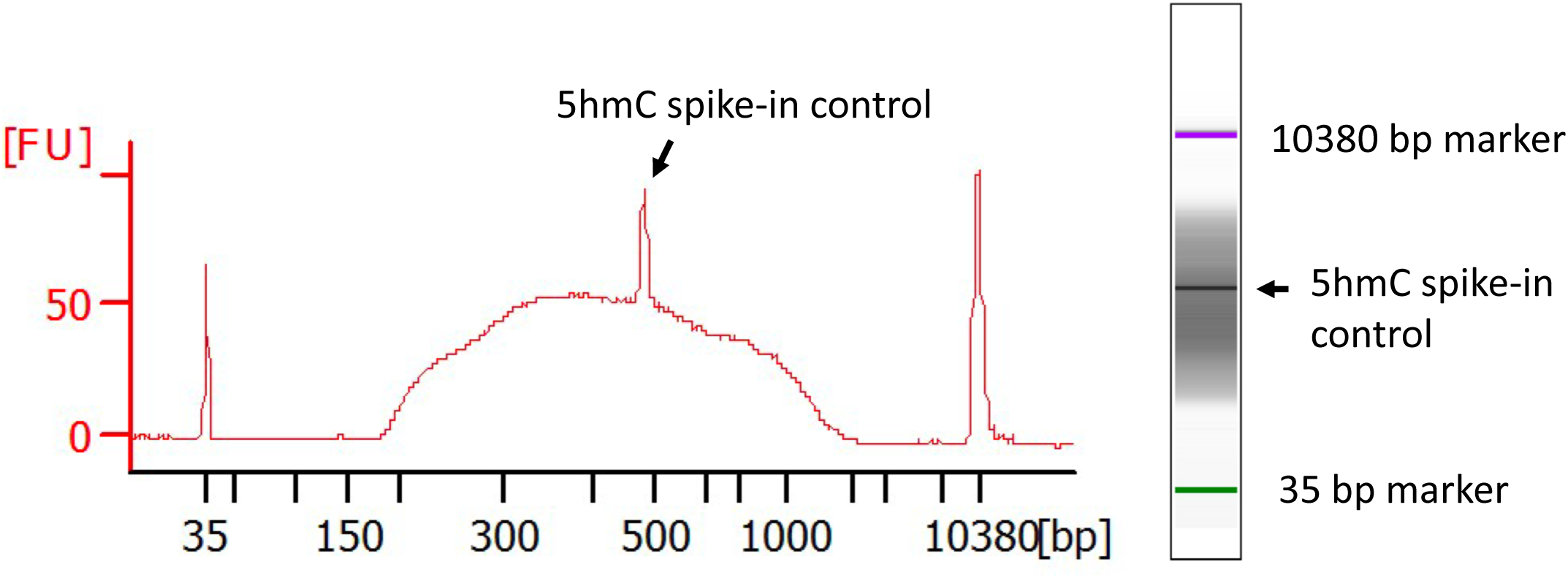
Agilent BioAnalyzer profile of a 5hmC Illumina library.

**Supplementary Figure 2:**
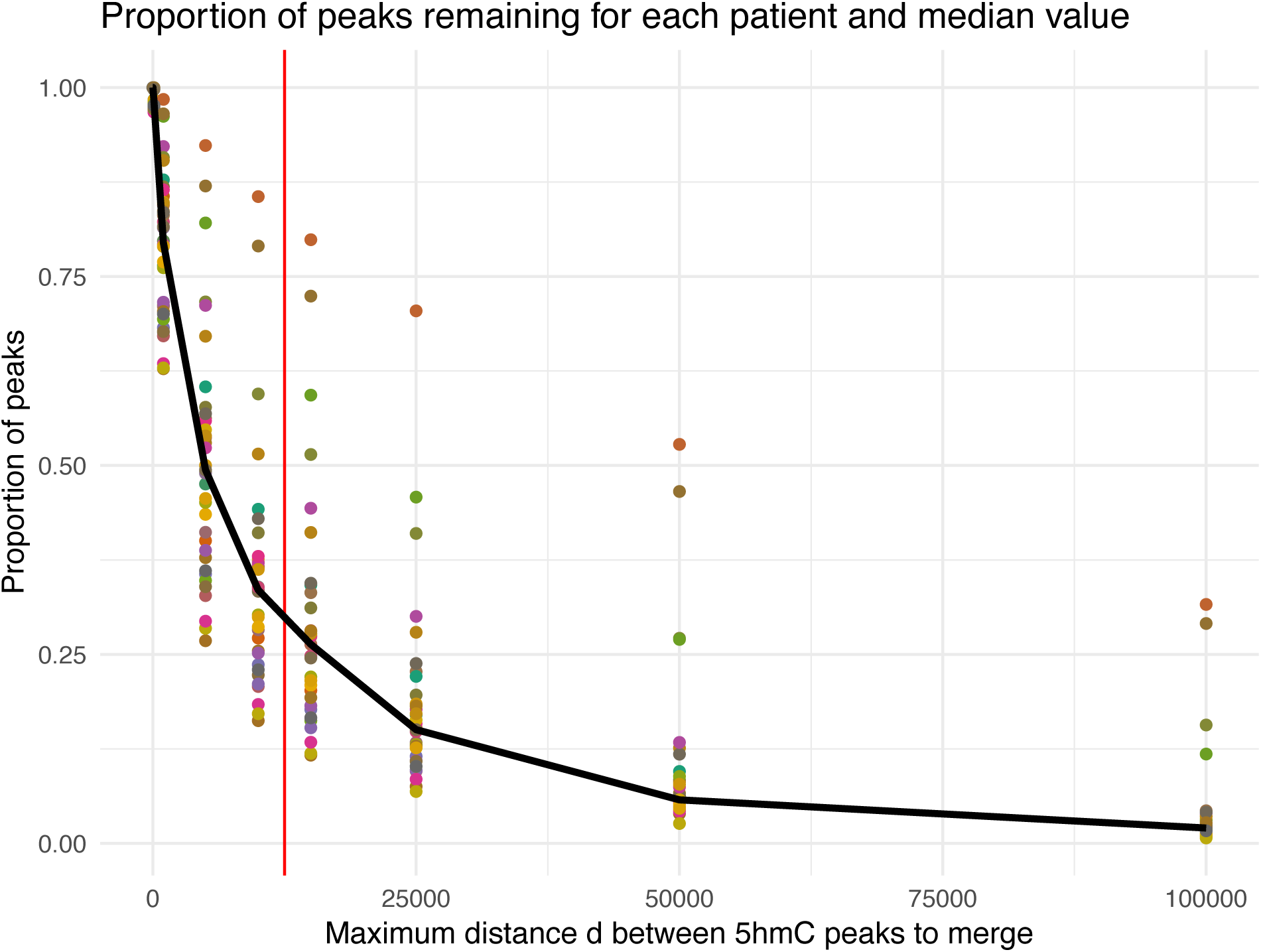
Criteria of 5hmC peaks to merge in 5hmC peak clusters. The y-axis represents the number of peaks left after merging. The x-axis represents the distance between peaks to merge. Each 5hmC samples were analyzed (one color per patient). The distance 12.5 kb was chosen to define the 5hmC peak clusters that we used in this study.

**Supplementary Figure 3:**
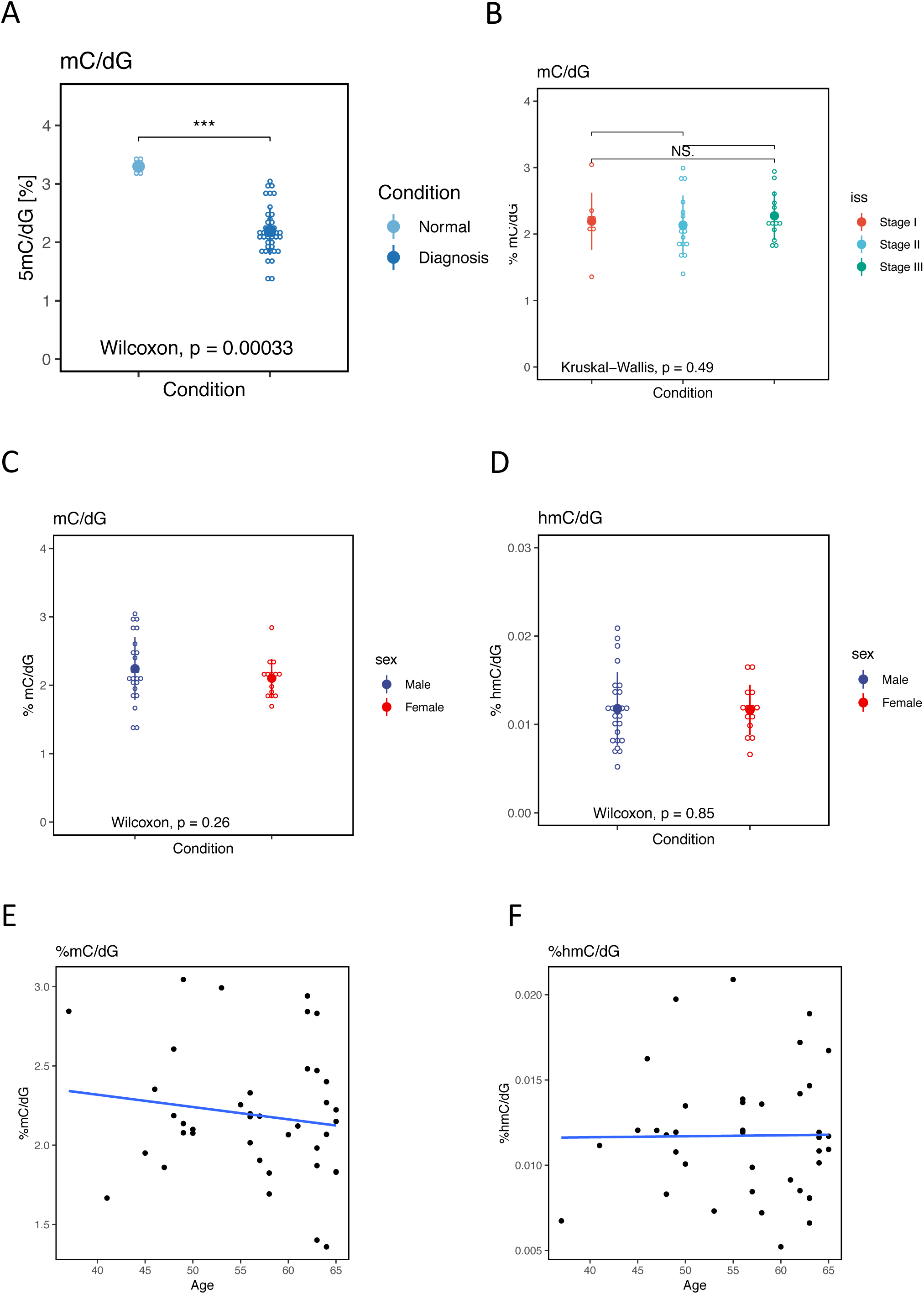
MS quantification of 5mC in genomic plasma cell DNA is independent of age and sex. (A) Dot plot of 5mC global quantification by MS in normal plasma cells from healthy donors (N=5), and of myeloma cells of patients at diagnosis (N=40). (B) Dot plot of 5hmC global quantification by MS by disease stage (ISS I N=9; ISS II N=17; ISS III N=13; NA=1). 5mC (C) and 5hmC (D) dot plot of MS quantification depending on the sex of the patients. 5mC (E) and 5hmC (F) dot plot of MS quantification depending on the age of the patients.

**Supplementary Figure 4:**
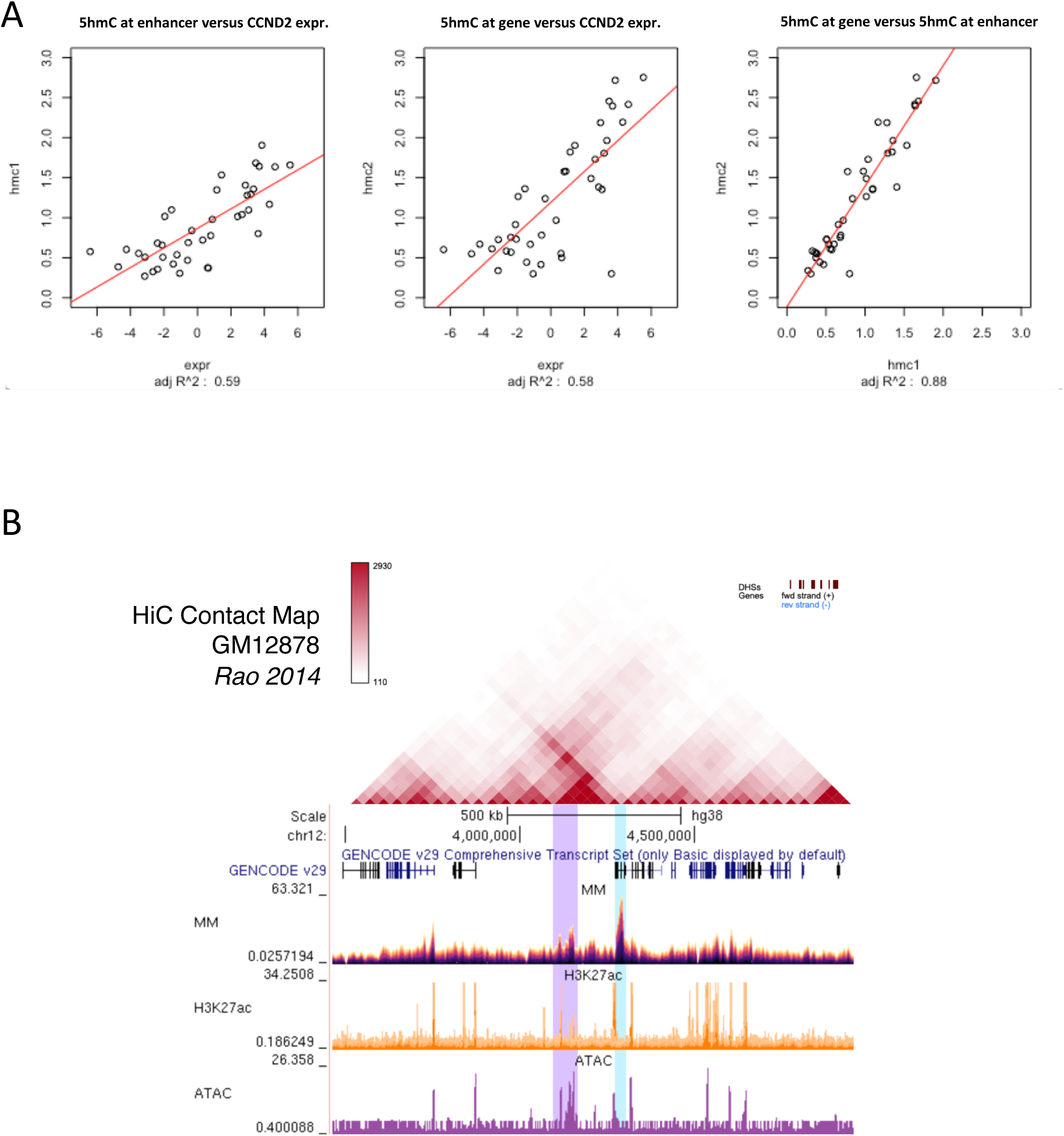
5hmC allows the identification of a putative CCND2 enhancer. (A) Correlation between *CCND2* expression, 5hmC at *CCND2* gene body and 5hmC at the putative 5hmC enhancer across the 40 MM patients. (B) Hi-C signal in lymphoblastoid cells (GM12878 cells) at the *CCDN2* locus showing the spatial interaction between *CCND2* gene and its putative enhancer.

**Supplementary Figure 5:**
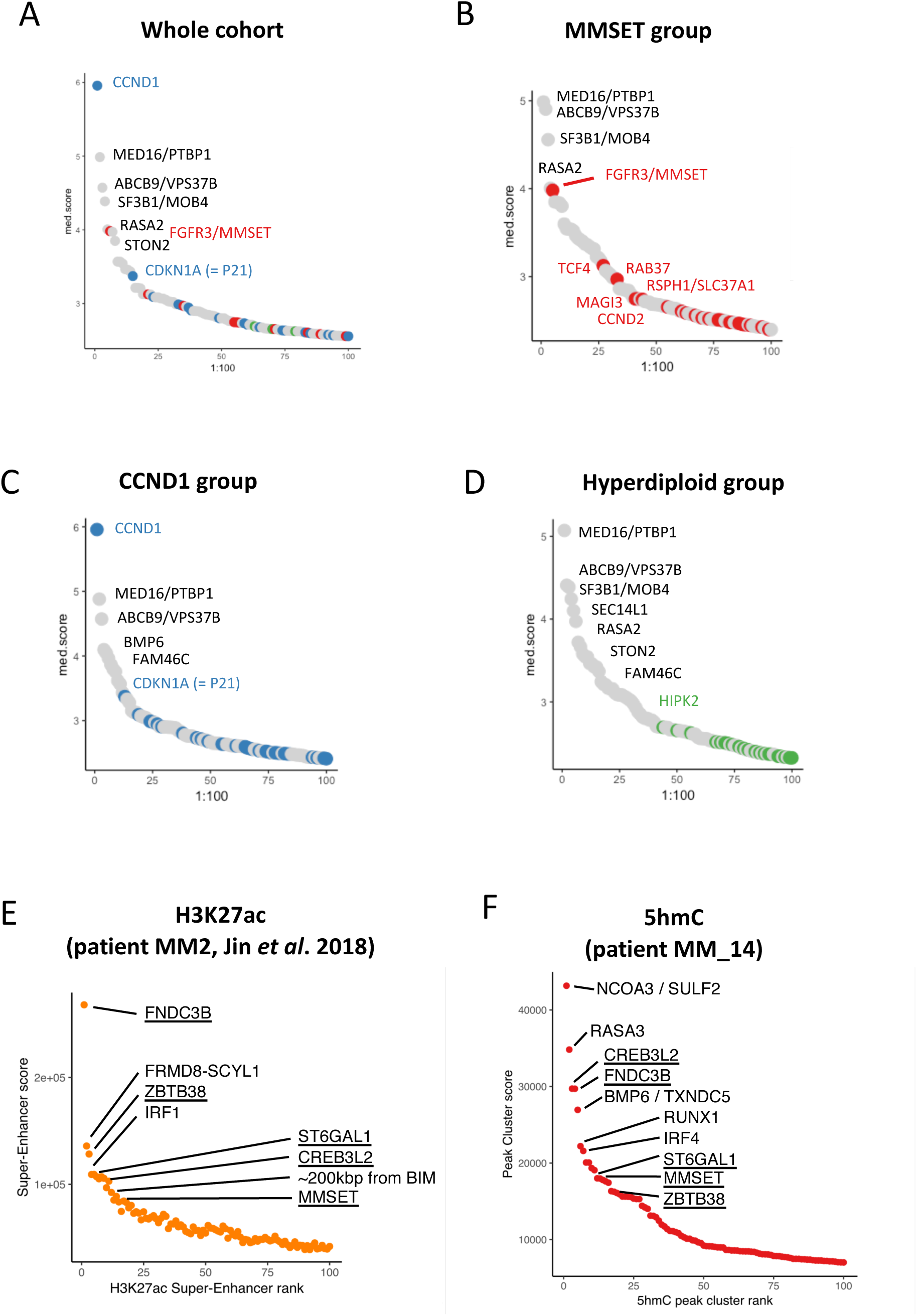
MM 5hmC peak clusters associate with H3K27ac super-enhancers. Rank ordering of the 100 strongest 5hmC peak clusters in the cohort (A), in the MMSET group (B), in the CCND1 group (C) and in the hyperdiploid group (D). (E) Rank ordering of the 100 strongest super-enhancers based on the H3K27ac signal in a MM patient with genomic anomalies t(4;14)/del13/amp1q (patient MM2 from Jin and colleagues^39^). (F) Rank ordering of the 100 strongest 5hmC clusters of a MM patient with genomic anomalies t(4;14)/del13/amp1q (our study: patient MM_14).

**Supplementary Figure 6:**
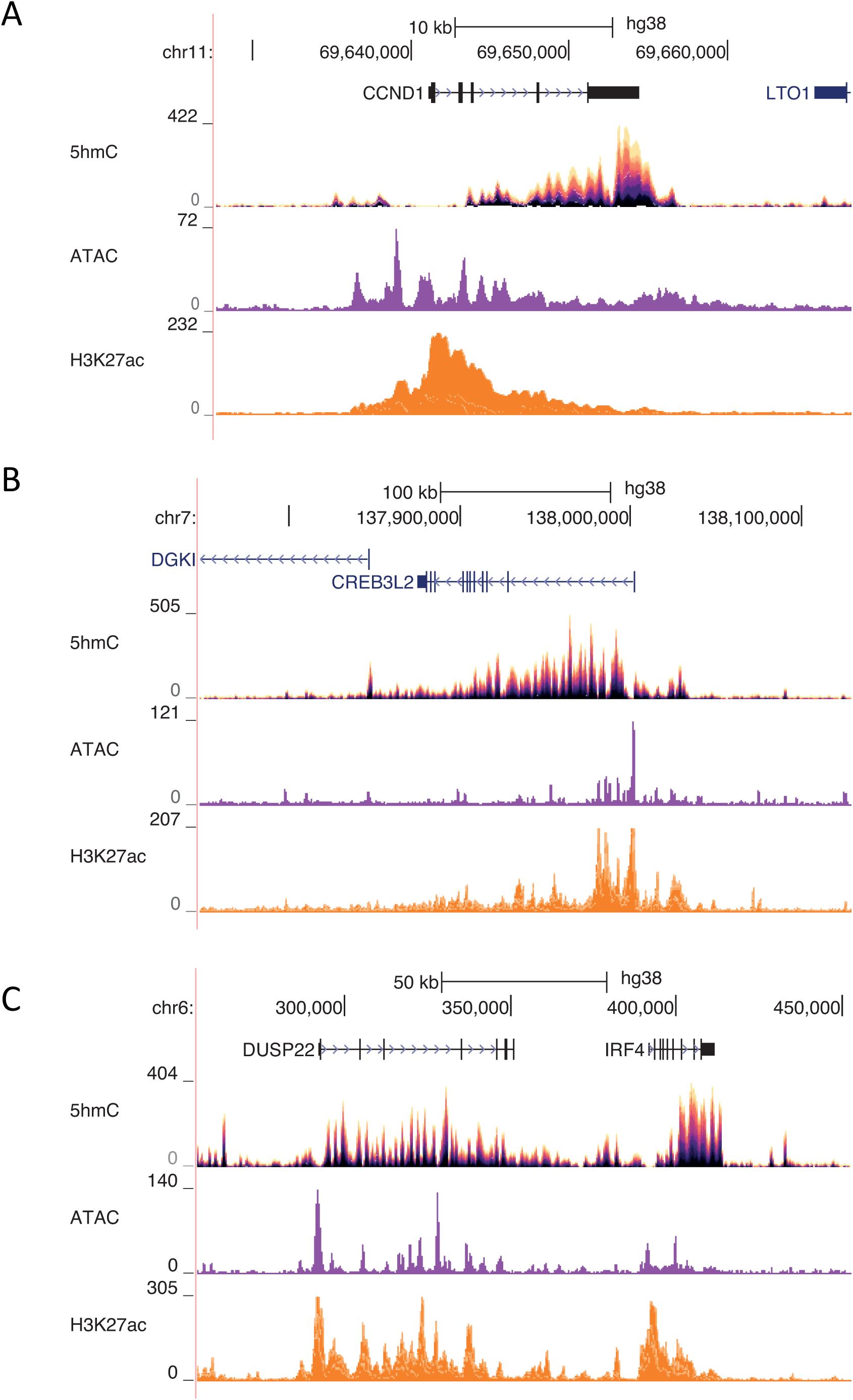
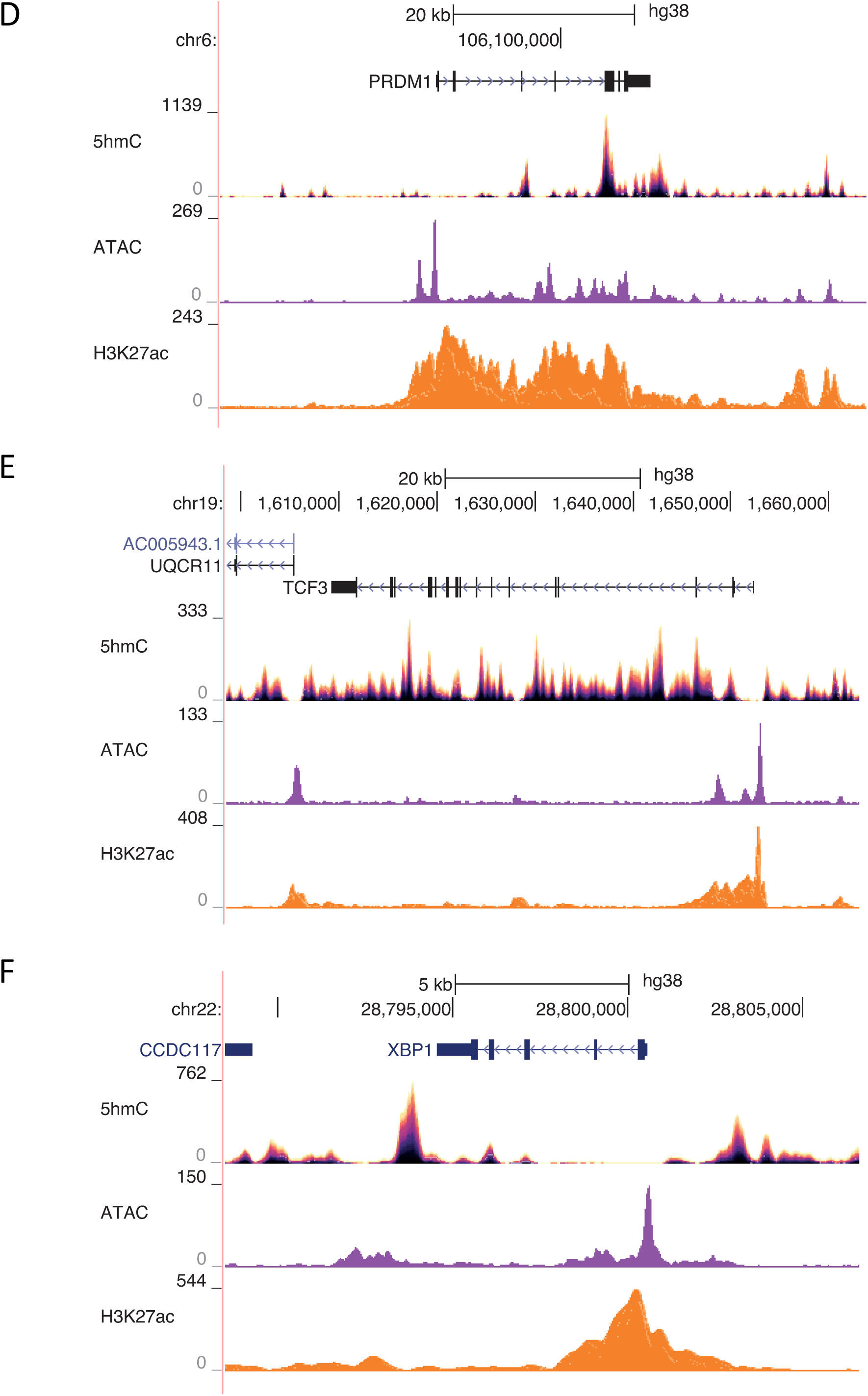
5hmC, ATAC and H3K27ac are enriched at major MM genes. Signal tracks of 5hmC, ATAC and H3K27ac at the loci *CCND1* (A), *CREB3L2* (B), *DUSP22* /*IRF4* (C), *PRDM1* (D), *TCF3* and *XBP1*.

**Supplementary Figure 7:**
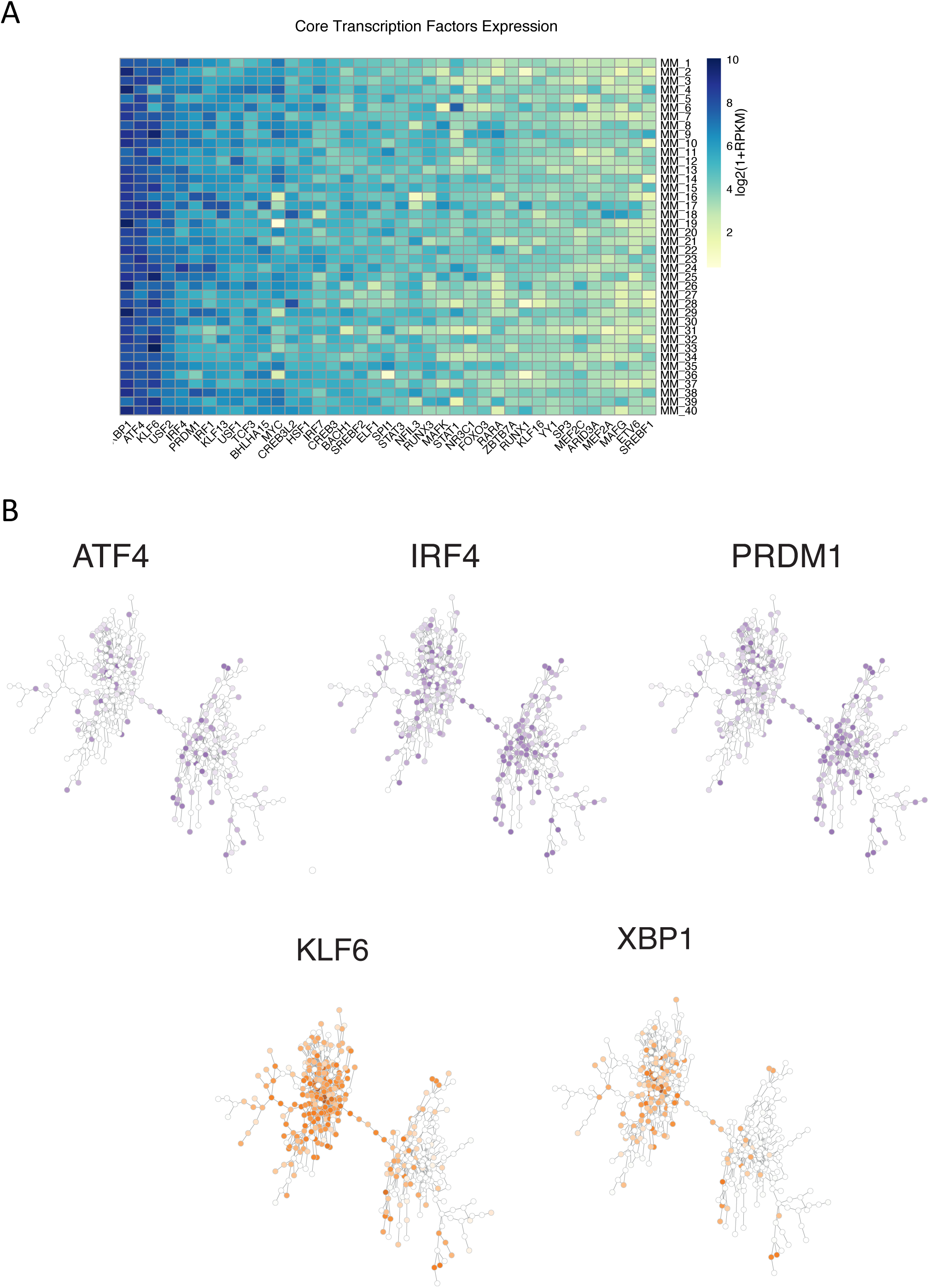
Core transcription factors binding motifs spread over the two networks. (A) Heatmap representing the expression level of the core transcription factors in the MM cohort. (B) Network representation of the 5hmC peak clusters linked with the strongest correlations. Each dot is a 5hmC peak cluster. Edges are represented if the correlation between is 2 nodes is greater than 0,715. Color intensity represents the normalized number of motifs ATF4, KLF6, IRF4, PRDM1 and XBP1 in the 5hmC peak clusters.

**Supplementary Figure 8:**
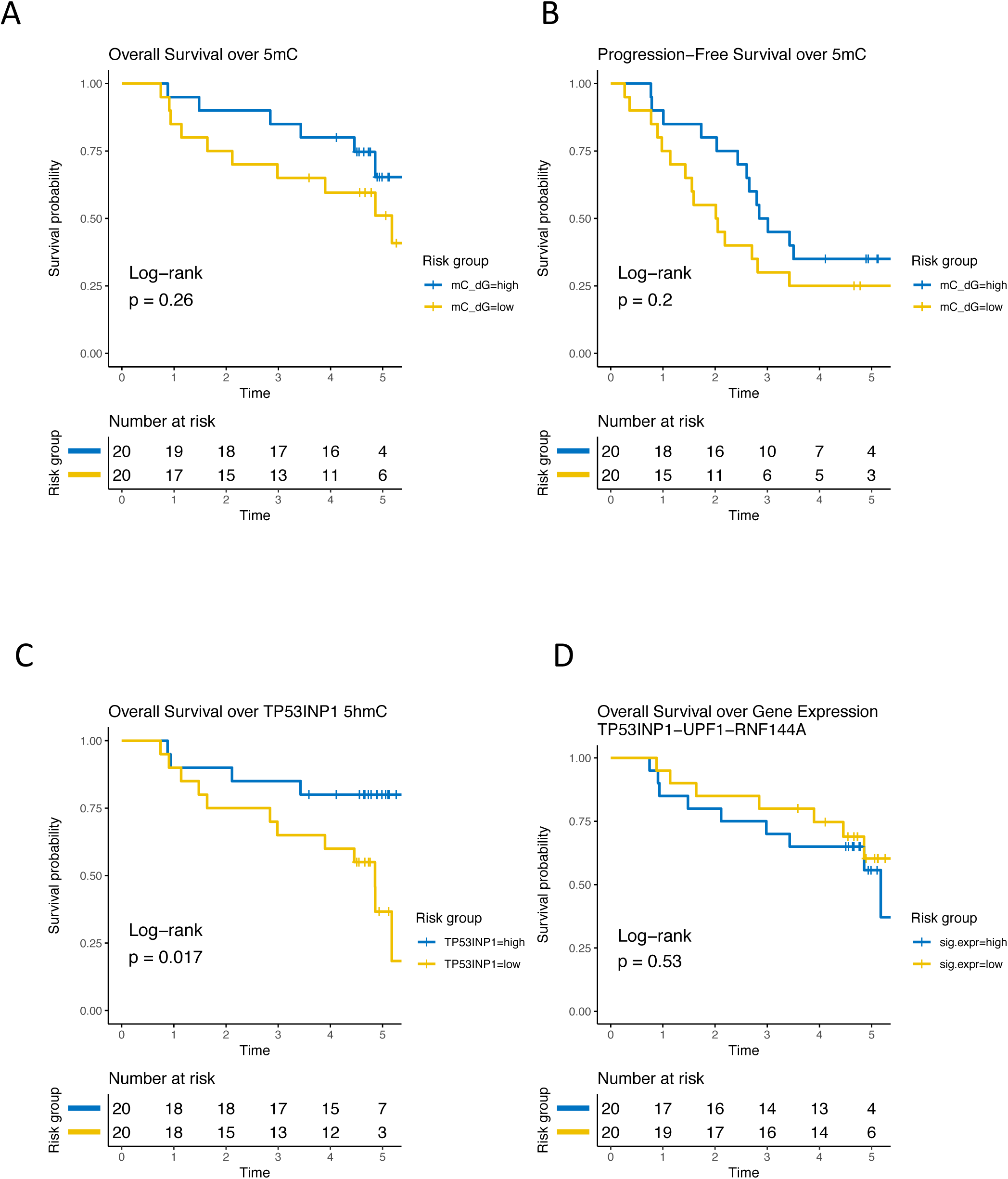
Survival courses depending on 5mC and 5hmC. (A) OS over global level of 5mC in MS. (B) PFS over global level of 5mC in MS. (C) OS over 5hmC enrichment at *TP53INP1* genomic locus. (D) OS over the score of the signature of *TP53INP1, UPF1* and *RNF144A* RNA expression levels (cf Methods). (Time: Number of years).

### Supplementary Table

**Supplementary Table: Description of the patients and genomic regions of interest.** (Tab1) Patients samples information. (Tab2) Principal Component Analysis. (Tab3) Group-specific regions. (Tab4) TF network analysis. (Tab5) Diagnosis/Relapse analysis. (Tab6) Survival analysis.

